# What do we gain when tolerating loss? The information bottleneck wrings out recombination

**DOI:** 10.1101/2021.08.27.457981

**Authors:** Apurva Narechania, Dean Bobo, Rob DeSalle, Barun Mathema, Barry Kreiswirth, Paul J. Planet

**Affiliations:** Institute for Comparative Genomics, American Museum of Natural History, New York, NY; Section for Hologenomics, The Globe Institute, University of Copenhagen, Copenhagen, Denmark; Department of Ecology, Evolution, and Environmental Biology, Columbia University, New York, NY, USA; Department of Epidemiology, Mailman School of Public Health, Columbia University, New York, NY; Center for Discovery and Innovation, Hackensack Meridien Health, Nutley, NJ; Division of Infectious Diseases, Children’s Hospital of Philadelphia, Philadelphia, PA; Perelman School of Medicine, University of Pennsylvania, Philadelphia, Pennsylvania, United States of America

## Abstract

Most microbes have the capacity to acquire genetic material from their environment. Recombination of foreign DNA yields genomes that are, at least in part, incongruent with the vertical history of their species. Dominant approaches for detecting these transfers are phylogenetic, requiring a painstaking series of analyses including alignment and phylogenetic tree reconstruction. These traditional pan-genomic methods do not scale. Here we propose an unsupervised, alignment-free and tree-free technique based on the sequential information bottleneck (SIB), an optimization procedure designed to extract some portion of relevant information from one random variable conditioned on another. In our case, this joint probability distribution tabulates occurrence counts of k-mers against their genomes of origin with the expectation that recombination will create a strong signal that unifies certain sets of co-occuring k-mers. We conceptualize the technique as a rate-distortion problem, measuring distortion in the relevance information as k-mers are compressed into clusters based on their co-occurrence in the source genomes. The result is fast, model-free, lossy compression of k-mers into groups that learns tracts of shared genome sequence differentiating recombined elements from the vertically inherited core. We show that the technique yields a new recombination measure based purely on information, divorced from any biases and limitations inherent to alignment and phylogeny.

**Significance:** The information bottleneck, a lossy compression technique borrowed from the information theoretic and Natural Langauge Processing literature, is well suited to detecting evolutionary patterns in sets of co-occuring k-mers. Here we show that we can detect simulated and real recombination events while highlighting a core set of k-mers that comprise the vertically inherited portion of any set of genomes. Moreover, the compressibility of any given set of genomes offers a new way to compare the pangenomes of clades across the microbial tree of life. In our application, the bottleneck is informed by genome origin, our relevance variable, but the technique is general. The information bottleneck can be used for any biological contingency matrix where the goal is to learn groups from unstructured data.

## Introduction

Microbial genomes have accumulated at an unprecedented rate^1,2^. Most work on their molecular evolution is grounded in sequence alignment and phylogenetic tree reconstruction, computationally expensive techniques ill-equipped to handle the volume of sequence encountered in a read streaming era^3^. The evolution of microbes is particularly challenging because recombined elements contribute signal unrelated to vertical descent^4^. Current techniques require alignment of reads across a reference genome^5,6^, whole genome alignment^7^, and/or phylogenomic methods based in orthology^8–10^. Each requires careful curation and deliberate sampling to limit data to reasonable scales. For larger, unbiased datasets that include as much natural variation as possible, these approaches are unsustainable. We need tools that can tolerate information loss without sacrificing knowledge of key evolutionary events.

Lossy compression, where an individual or algorithm makes decisions about which data are important (or relevant) from a large body of information, may offer a solution^11^. To do this in a principled way, the relevance of a given dataset can be measured as information retained about some other correlated variable. For example, in unsupervised natural language processing (NLP) large corpora of texts are distilled to a few topics that reflect overall themes by comparing patterns of co-occurring words in the source texts. In topic modeling of this sort, the texts themselves are the relevance variable. The goal is to cluster the word distribution with respect to the documents from which they arise. This idea was first described by Tishby, Pereira and Bialek as the information bottleneck (IB)^12^. It was premised on rate distortion, Shannon’s original theory of lossy compression which yoked signal distortion to the rate at which that signal can be encoded^13^. The IB’s primary innovation was the use of a relevance variable to quantify this distortion and instill it with meaning. Topic modeling was one of this technique’s first applications.

Topic modeling has become an important part of the NLP literature with a number of wider applications to unsupervised machine learning. The dominant technique in the field is Latent Dirchilet Allocation (LDA)^14^, a probabilistic method, that like the IB, considers each document a mixture of topics. Some groups have applied this idea to whole genomes^15–17^, and since the publication of STRUCTURE, LDA has become foundational in the genetics literature^18^. Despite LDA’s popularity and success, a number of authors have shown that unbalanced sampling can lead to erroneous or missed population assignments^19^. LDA also makes a number of statistical assumptions including the assignment of hyperparameters and a Dirchilet prior^20^. In contrast, the IB is model free, unaffected by size sample bias.

Genomes are living documents that can be sliced into words of arbitrary size^21,22^. In a genomic context, where words are k-mers and documents are their genomes of origin we hypothesize that IB derived topics could represent co-occurring groups of k-mers that highlight shared ancestry. K-mers might be arranged in co-linear blocks, distributed across the genome as fragments inherited simultaneously, or may be common to all genomes, providing a simple, operational definition of a genomic core. In all cases, lossy compression of k-mers into topics is guided by how often they co-occur with respect to their genomes of origin. The process requires only two parameters: the k-mer size and the number of clusters expected. In the NLP topic modeling analogy, the core genome of a species could be considered the set of concepts common to every document in a library, while recombined regions are like themes or ideas restricted only to certain shelves.

Here we apply the IB to tens of thousands of microbial genomes. Remarkably, our approach identifies recombination tracts without making any attempt to model evolution, annotate genes, build alignments, or reconstruct trees. Further, squeezing k-mers through the bottleneck yields a kind of distortion we repurpose to measure horizontal signal.

## Theory and Implementation

Consider a set of genomes (Y) chopped into k-mers (X). The information bottleneck is designed to compress X under the constraint of Y^12^. The IB extends Shannon’s theory of rate distortion by guiding it with an additional, orienting variable. In our case, this variable is genomes of origin. This concept of a relevance variable assigns value to the resulting distortion. The choice of Y defines relevant features in the signal. If X and Y are tabulated as a joint probability distribution, the information that X provides about Y is squeezed through a simpler representation to learn a set of T groups (Figure 1). In a rate distortion sense, T can be considered the channel capacity. For the technique to work, the two variables in our joint distribution *p(x,y)* must be non-independent, or more precisely, must have positive mutual information, *I(X,Y)*:

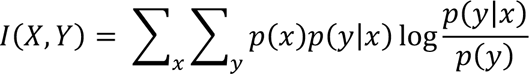

**Figure 1.**
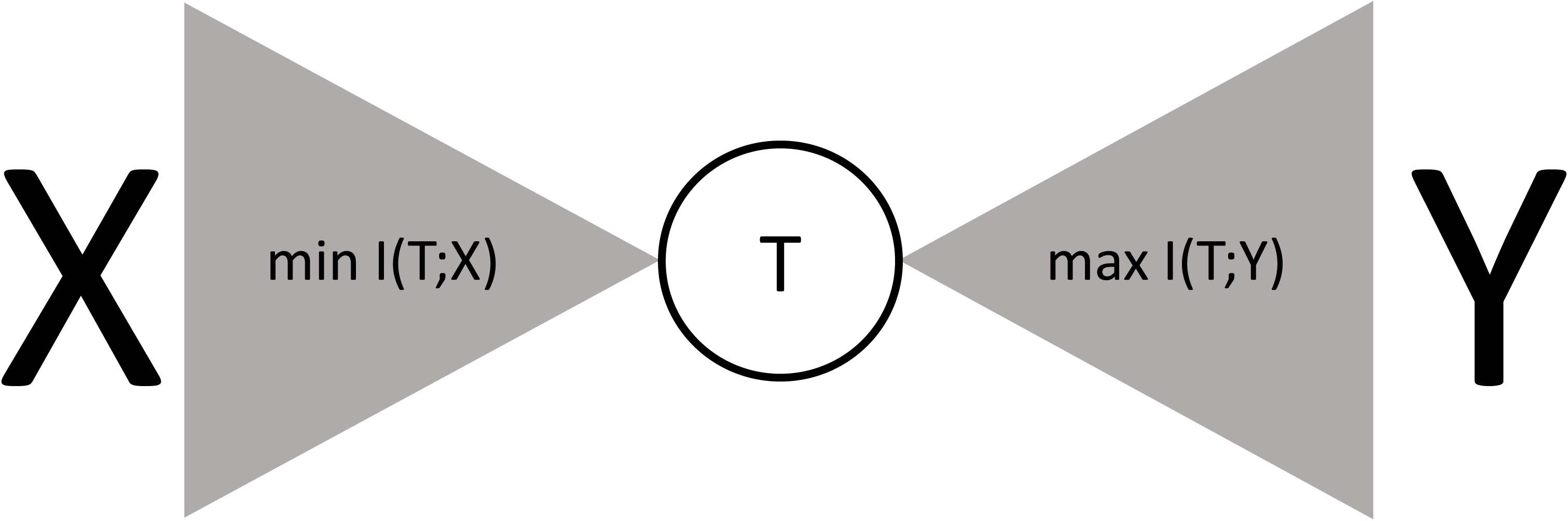
The information bottleneck. In the information bottleneck a distribution, *X*, is compressed into *T* while retaining as much information as possible about a correlated relevance variable, *Y*. The joint distribution, *p(x,y)*, has positive mutual information and the goal of the information bottleneck is to capture as much of that information as possible at interpretive scale. The technique is a classic optimization problem wherein the mutual information between *T* and *X* is minimized, while the mutual information between *T* and *Y* is maximized. At optimality, *T* is presumed to be a lossy but adequate model of *X*.

T is now a meaningful compression of the data, maximizing the mutual information between the clusters and genomes, I(T;Y), while minimizing the mutual information between the k-mers and the clusters, I(T;X). The IB is a classic optimization problem (Figure 1).

Compressing X into T is equivalent to minimizing the following Lagranian with respect to the source genomes, Y:

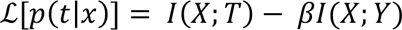

This formulation balances the compactness of X, with the erosion of information about Y. β is a multiplier that slides through the optimization landscape. As beta approaches 0, k-mers are clumped into fewer and fewer clusters, emphasizing compression. As beta approaches infinity, every k-mer is its own cluster, preserving all relevant information. Of course, collapsing all k-mers into one cluster is overly reductive, and assigning each k-mer to its own cluster is meaningless. The IB negotiates these two extremes. In NLP, the result is a set of clusters that coalesce into topics over a body of literature^23^. In genomics, the process identifies co-occurring and/or spatially co-located k-mers with distinct biological and/or evolutionary meaning.

Minimizing the Lagranian above has an exact, optimal solution. The most surprising outcome of this solution is that the relative entropy, or Kullback Liebler divergence^24^, emerges as the distortion measure for the information bottleneck. The relative entropy is a fundamental quantity in information theory, and in our IB context, it measures the distortion of compressed k-mers:

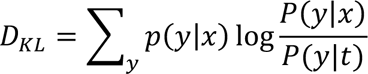

We set the number of clusters and implement a sequential clustering procedure. From an initial random distribution of all k-mers across our clusters we draw one k-mer out, and represent it as a singleton. Using greedy optimization, we merge this singleton into one of the existing bulk clusters. Slonim’s sequential information bottleneck (SIB)^25^ employs the Jensen-Shannon divergence^26^ in the cost of merging a k-mer, x, into a cluster, t:

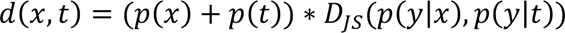

A k-mer will join a new cluster only if its new address reduces the total distortion. Otherwise, it remains in its existing cluster. With respect to our initial random conditions, this algorithm is guaranteed to converge to a local optimum. We mitigate the risk of getting trapped in local optima by testing several random initializations, and we reduce the overall computation by implementing a k-mer sketch^27,28^.

Once the clusters stabilize, we quantify the information captured by calculating the normalized mutual information, *NMI = I(T;X) / I(X;Y).* Trivially, the NMI is one when each k-mer occupies its own cluster and is zero when all k-mers are mixed into one cluster. The curve traced between T = 1 and T = *x* is called the relevance compression curve^29^. The choice of T is analogous to the optimization of β in the Lagranian above. Both modulate the Shannon channel capacity, but we vary T for the deterministic case involving hard clustering. As with β, the shape of this curve describes the compressibility of the data.

The most important aspect of the SIB, and the reason we chose it for this work, is that its distortion, as measured by NMI, is a proxy for horizontal signal.

## Results and Discussion

### The bottleneck in test: simulated recombination events

We simulated 50 recombination events across a set of 100 1 Mbase genomes in SimBac^30^. These events appear in black on the innermost track if the circos plot in Figure 2. We sketched 19-mers from all 100 genomes and applied the information bottleneck, modeling 50 clusters. The outermost track is a frequency plot of k-mers from the core cluster aligned to arbitrary genome from the set. The core emerges as a dense block of shared genome sequence interrupted by our simulated recombination events. K-mers that would otherwise occupy these gaps are sorted into other clusters. Because the compression is driven by genome of origin, if a single ancestor sustains multiple transfer events, all k-mers from these events merge into a single cluster shared by the same subset of descendants. Plots of the bottleneck-defined core function almost as a photographic negative, highlighting the blank spaces as regions scrambled by horizontal signal.

**Figure 2.**
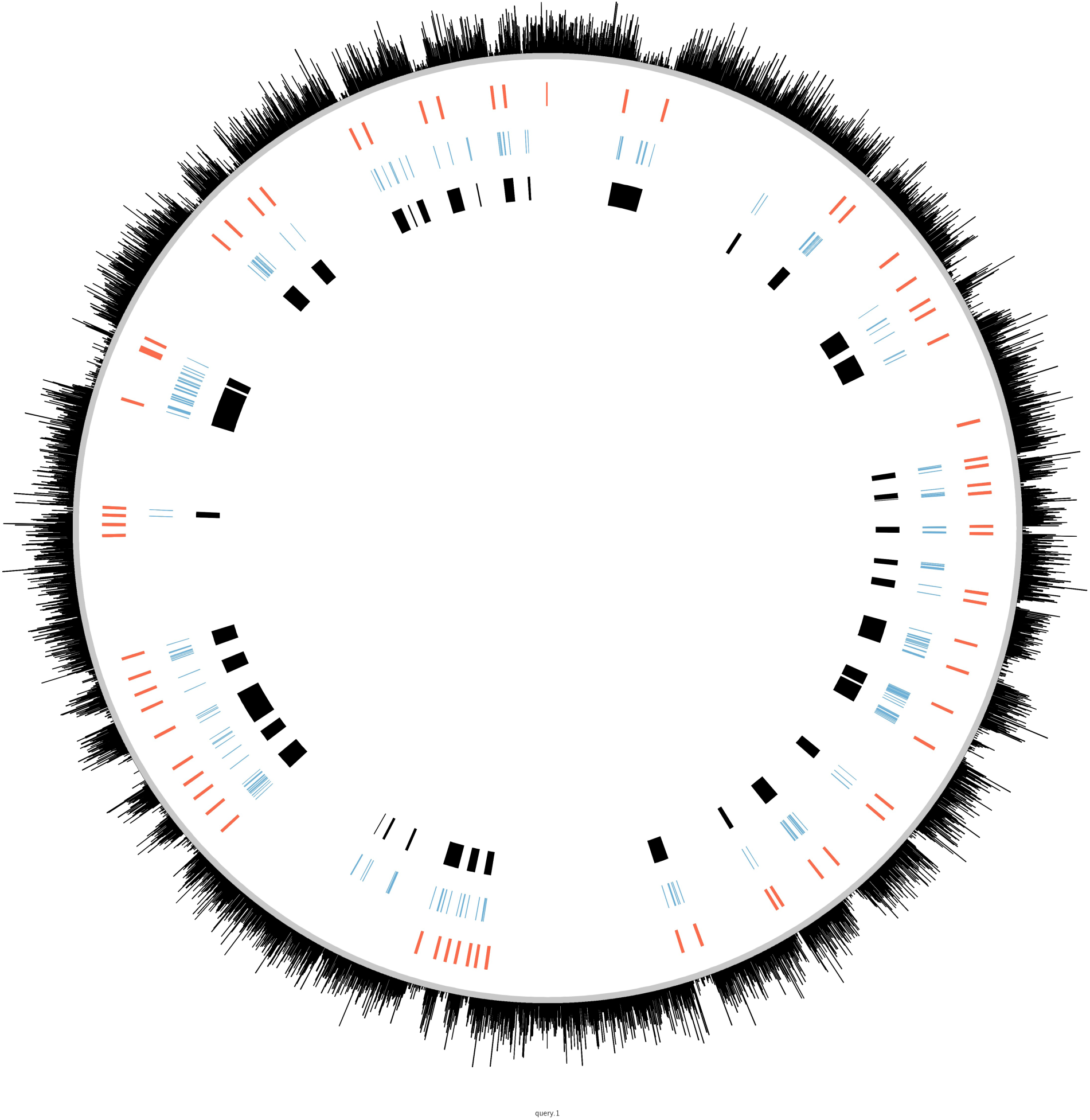
Simulations. We simulated 50 recombination events across 100 one megabase genomes. The recombination events are shown as black blocks on the innermost track. IB based changepoint calls for breakpoints are shown in red, and ClonalFrameML based intervals are shown in blue.

Meaning emerges from this spatial organization of k-mers and corresponds to regions of shared ancestry. In the examples shown, the pattern of events is evident by eye, but with increasing input genomes and increased modeling (more clusters), automated inspection is key. We use a signal processing method based in change point detection to programmatically detect changes in k-mer frequency. We specifically employ the PELT algorithm^31^ to model probabilities of change along the genome backbone. As shown in the track of red hashes, changepoint detection largely delimits the boundaries of our simulated recombination events. PELT’s detection sensitivity is modulated by a penalty parameter that can vastly change the analysis. A sweep through values for this parameter is shown in Supplementary Figure 1. While this parameter can alter the changepoints derived from our primary frequency data, the frequency data itself almost always shows gaps in the core’s continuity at recombination events. In the example shown in Figure 2, 48 of the 50 recombination events have k-mer counts significantly lower than the rest of the core’s background (Wilcoxon, p < 0.05).

We compare our approach to ClonalFrameML^6^, the leading recombination detection program in microbial genomics. Unlike the IB, ClonalFrameML requires a reference, alignment of all remaining 99 genomes to that reference, tree reconstruction from the resulting character matrix, and recombination detection using a likelihood-based sliding window approach. ClonalFrameML calls are shown in the track of light blue hashes. The program successfully identifies regions of recombination, even picking up smaller regions that IB fails to detect at the shown change point sensitivity. But the program seems to subdivide events into smaller ones, sometimes up to 20 smaller events in one large expanse.

Lossy compression drops data. Forcing all the signal in our k-mer occurrence matrix through a narrow 50 cluster channel garbles portions of the original message. This is equivalent to channel noise. It recalls Shannon’s original formulation of the rate distortion problem^13^. In this case, modeling more clusters increases the rate of transmission, and reduces the distortion of the message received. With respect to the information bottleneck, we quantify this effect using a relevance-compression curve^32^. In Supplemental Figure 2, we show data from 100 simulated sets each of 4, 10, 100 and 1000 one megabase genomes. Regardless of the number of genomes in each experiment, the NMI increases with increased modeling (one example for each shown in Panel A). As expected, this increase is rapid and stark in the smaller datasets; slow and somewhat halting in the larger ones. Complex scenarios with more genomes and more evolutionary paths require a broader channel (more clusters) to transmit signal. The theoretical extremes for the relevance-compression curves are intuitive. At the origin, all the relevant information is destroyed. With too many clusters, we retain more information than we can interpret. The curve traced between these two extremes is a fingerprint of the data. A convex shape suggests natural structure easily modeled with just a few clusters. Data that resists compression flattens this curve. Theoretically, the space above the curve is unachievable by any process, forming an upper bound. The relevance-compression curve therefore defines absolute limits on the quantity and quality of evolutionary information communicated as we sweep through a dilating channel. In Supplemental Figures 2B and 2C, we show how information loss in more complex compression problems is driven by the narrowing gap between k-mer frequency outside versus inside the simulated recombination events. With four genomes, discrimination is easy. With 1000, noise limits our resolution.

The relevance-compression curve’s responsiveness to evolutionary complexity opens the door to reconceptualizing recombination. Given a sequence set, most researchers would calculate R/theta^33^, the ratio of the recombination rate over the mutation rate, to quantify a species’ tendency to recombine. But R/theta’s calculation is grueling, requiring alignment (either reference based or across whole genomes), tree reconstruction, and algorithms to isolate the clonal frame^5,6^. The information bottleneck clearly pinpoints recombination events in space without resorting to any of these traditional procedures.

NMI is a corollary to this compression, but can NMI itself be an informative recombination metric? In Figure 3 we show that for sets of 100 simulated genomes, NMI decreases with increased recombination rate. We also show that even very high background mutation rates have a negligible effect on NMI for all sets except those subject to vanishingly low recombination. Recombination seems to increase distortion indicating that in our context evolutionary complexity is tied to information loss. At high rates of recombination, every base of a 1 Mb genome is likely scrambled. Under such flux, some sites recombine several times, eroding the core genome itself. This erosion is reflected in the steep decline in NMI with increasing recombination rate. The NMI’s responsiveness to both recombination rates and mutation rates makes it an attractive alternative to arduous – and in some cases – impossible R/theta calculations. As expected, we capture more information if we allow more clusters, but over-modeling in simpler evolutionary scenarios erodes the information captured (Supplemental Figure 3).

**Figure 3.**
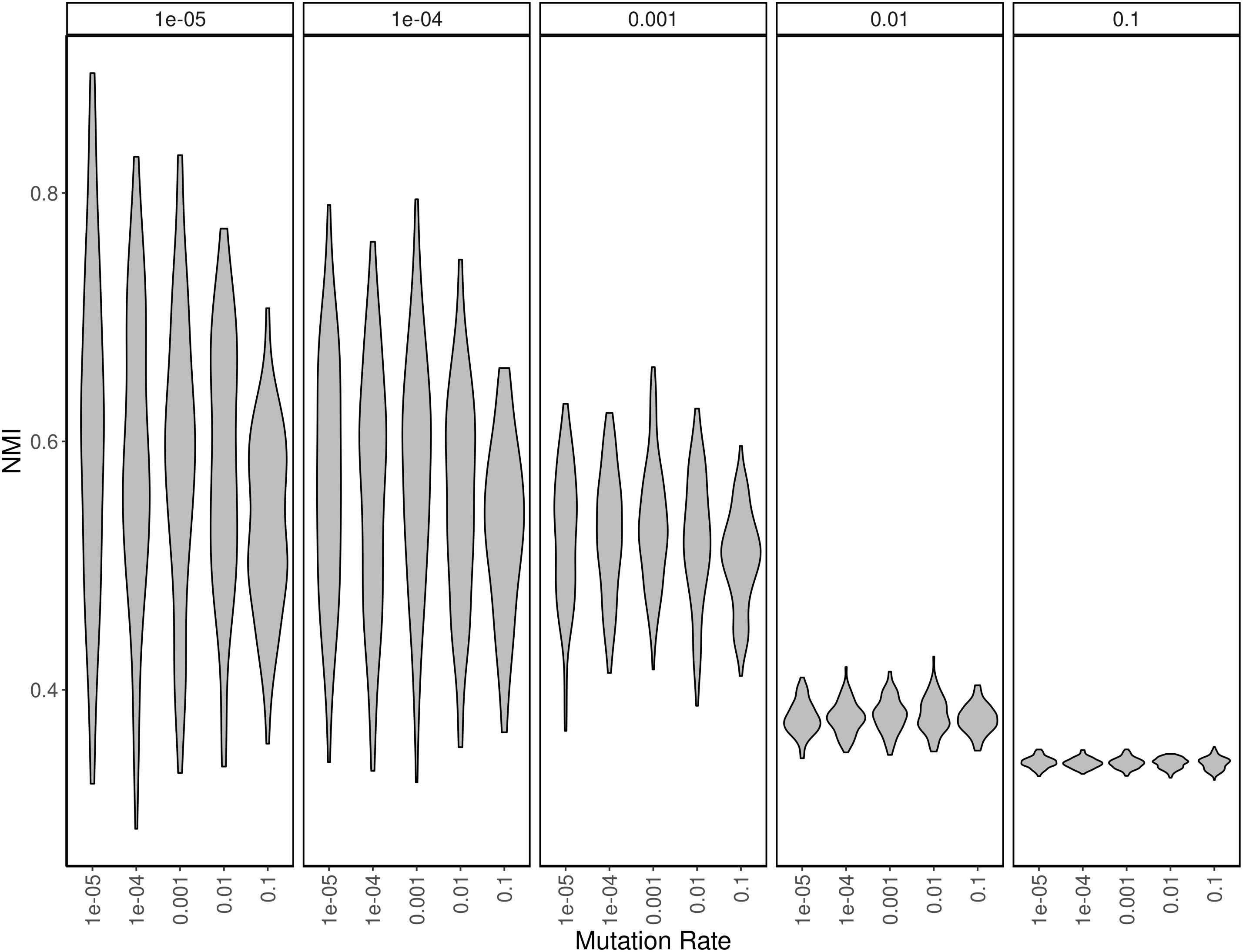
Simulation across various rates of recombination and background mutation. As the recombination rate increases, the NMI decreases. The effect of background mutation rate is swamped by recombination, becoming a factor only when recombination rates are very low.

### The bottleneck in action: real world recombination

We used genomes from ST239 *Staphylococcus aureus* to illustrate that our method can corroborate known, large scale recombination events found in nature. The ST239 strain is a hybrid: a segment from a CC30 (clonal complex 30) donor replaced nearly 20% of the homologous region in a CC8 strain^34^. The evolutionary histories of genes across these segments are incongruent. Previous studies compared the histories of thousands of genes to reach this conclusion^35^. Here, we attempt to localize this same phenomenon using the co-occurrence pattern of k-mers alone. We chose 10 genomes (GCA_000146385.1, GCA_000012045.1, GCA_000011505.1, GCA_000011265.1, GCA_000013425.1, GCA_000204665.1, GCA_000159535.2, GCA_000027045.1, GCA_000017085.1, and SA21300), sampled from both the donor clade (CC30), the recipient clade (CC8), and genomes outside of the evolutionary event.

Figure 4 highlights two of these 10 genomes, and three of the 60 clusters we modeled for this analysis. Both *S. aureus* COL (CC8) and *S. aureus* T0131 (ST239) share a large, congruent core. The gap in this core characterizes the dimensions of the recombination event, whose k-mers are split into two other clusters, shown here as the second and third tracks. The bottleneck learns the structural evolution of the clade as tracts of co-occurring sequence. The clusters themselves comprise an evolutionary model for the structural event and highlight the core. Genome origin guides k-mer compression by forming the basis of the distortion measure. We lose information in a controlled and quantitative way, and short circuit the intensive phylogenomic analyses required^35^ to identify the tract.

**Figure 4.**
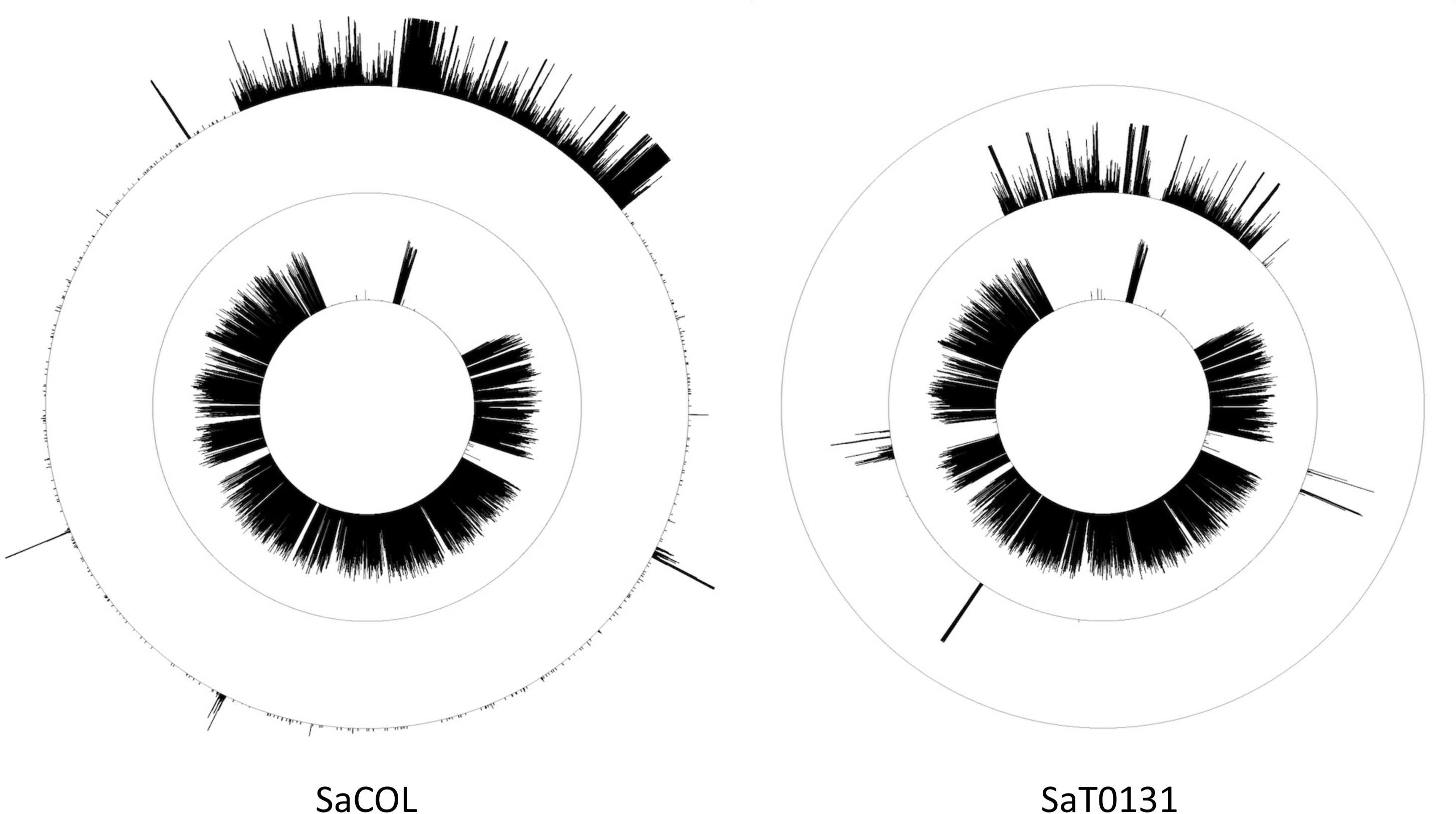
Modelling ST239’s hybridization event. We selected 10 *S. aureus* genomes to track the ST239 hybridization event with the information bottleneck. We chose COL to represent the CC30 donor strain, and T0131 the CC8 acceptor. Of the 60 clusters we calculated, we show the three that capture the hybridization event. The innermost track is a frequency plot of k-mers that define the core. The second and third tracks are flipsides of the HGT event that created ST239.

Our approach introduces a new type of comparative genomics based on compression. Figure 5 shows relevance compression curves for the ST239 genomes alongside 100 genomes of *Mycobacterium tuberculosis* and *Helicobacter pylori*. The convex shape of the *M tuberculosis* curve reflects its clonality. Data from a more open pangenome that resists compression flattens the curve. Highly recombinogenic species like *H. pylori* suffer this sort of steep information loss.

**Figure 5.**
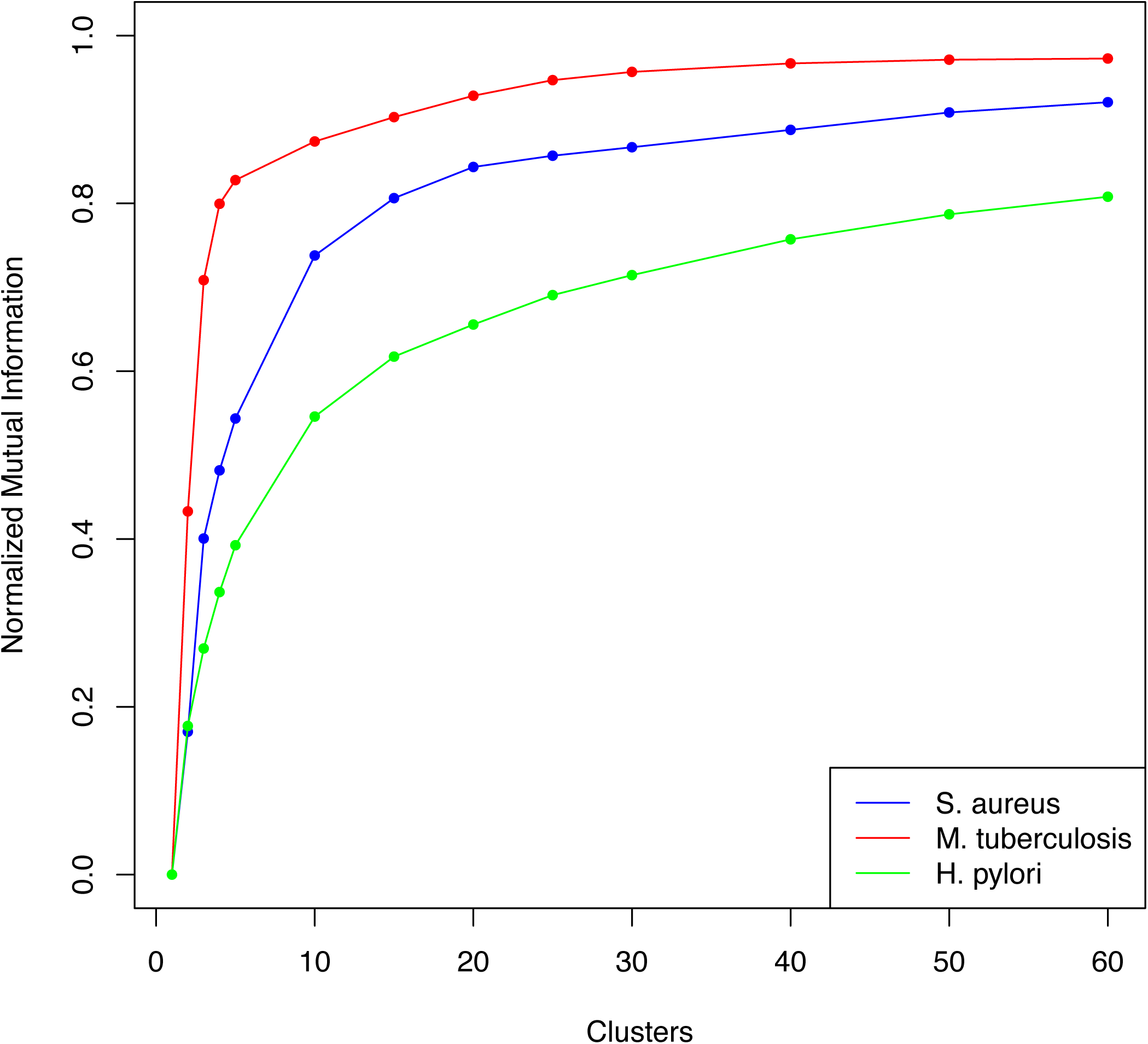
Relevance compression curves across three species. In an information bottleneck experiment, the relevance compression curve traces the increase in normalized mutual information with the number of clusters modeled. The curves quantify the amount of information lost at a given modeling threshold. We show how this type of relationship can function as a marker for evolutionary strategy by calculating curves for three very different groups of microbes: *M*. *tuberculosis*, a species thought to demonstrate little if any HGT; *S. aureus*, a species considered largely clonal with occasional HGT; and H. pylori, a species known to employ HGT as an engine for diversity (A).

In Supplementary Figure 4 we show relevance compression curves for the 28 most well studied microbial genomes in RefSeq. In all cases, increased modeling captures more information, but at varying rates. We also demonstrate that we can reach a steady-state NMI with a small number of randomly selected genomes regardless of clade (Supplementary Figure 5). We interpret the shape of the relevance compression curve as a proxy for evolutionary strategy and show that 50 randomly selected genomes can reflect the nature of the species.

### The bottleneck as a new measure of recombination

We have shown that in simulation, given a set number of clusters, increased recombination results in decreased NMI. To test whether this pattern holds for real data, we analyzed the top 13 most sequenced bacterial species in RefSeq with at least 100 complete, contiguous genome assemblies. For each of these 13 species, we randomly selected one complete genome as a reference, and 100 random genomes as queries. We calculated R/theta for these selections and repeated the experiment 100 times for all 13 species. Figure 6 shows R/theta (A) and r/m (B) plotted against the corresponding five cluster NMI. The data indicate that NMI is an inverse mirror of R/theta: higher levels of recombination tend to lower compressibility. NMI clearly offers a conceptual alternative to R/theta calculable in a fraction of the time (Figure 7). A simple distortion metric produced as sequence compresses through an information bottleneck provides a ready alternative to currently accepted approaches.

**Figure 6.**
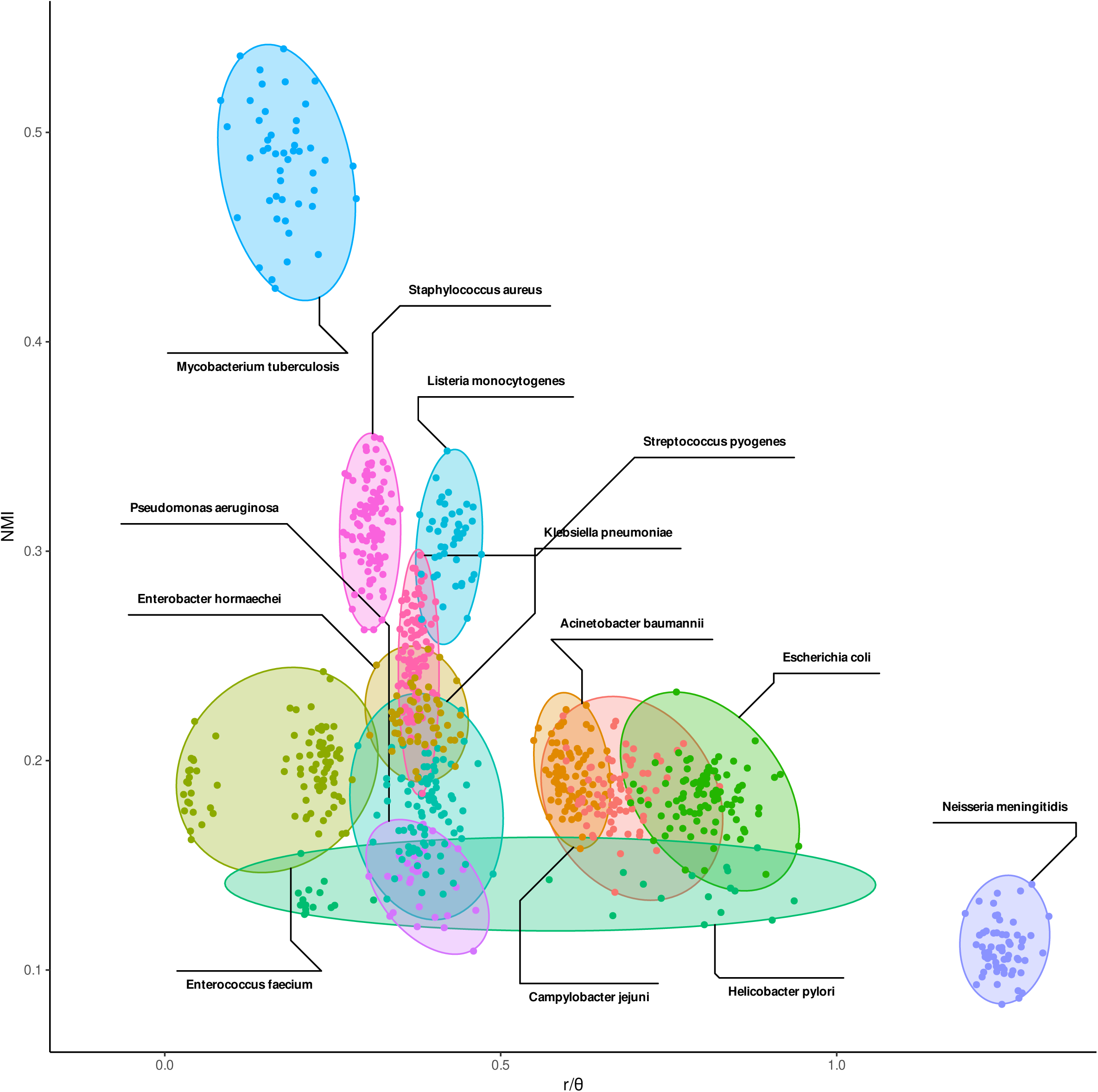
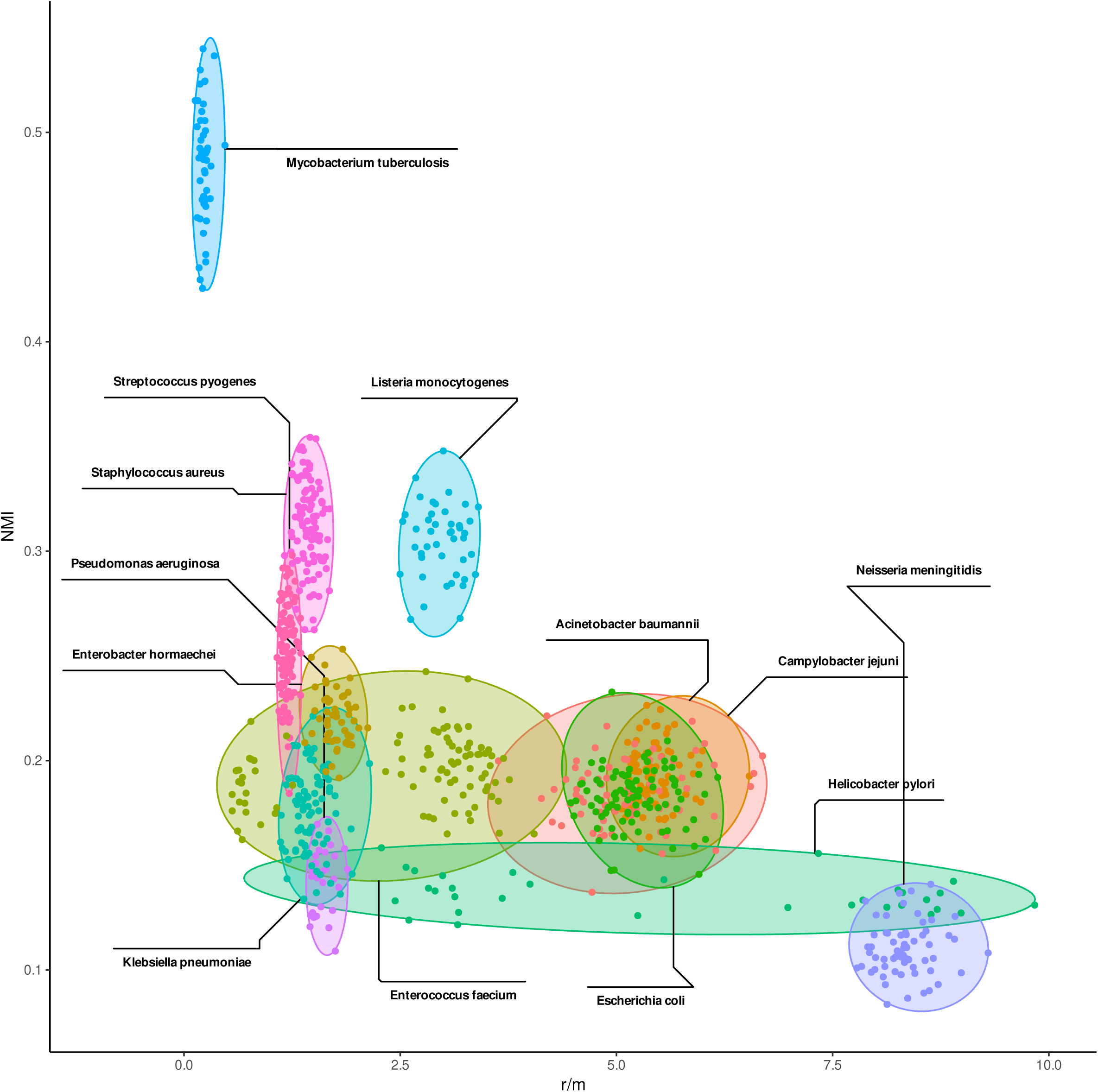
NMI tracks R/theta. We show that for 13 key species in Refseq, more recombinogenic species (as described by r/theta in panel A and r/m in panel B) are less easily compressed.

**Figure 7.**
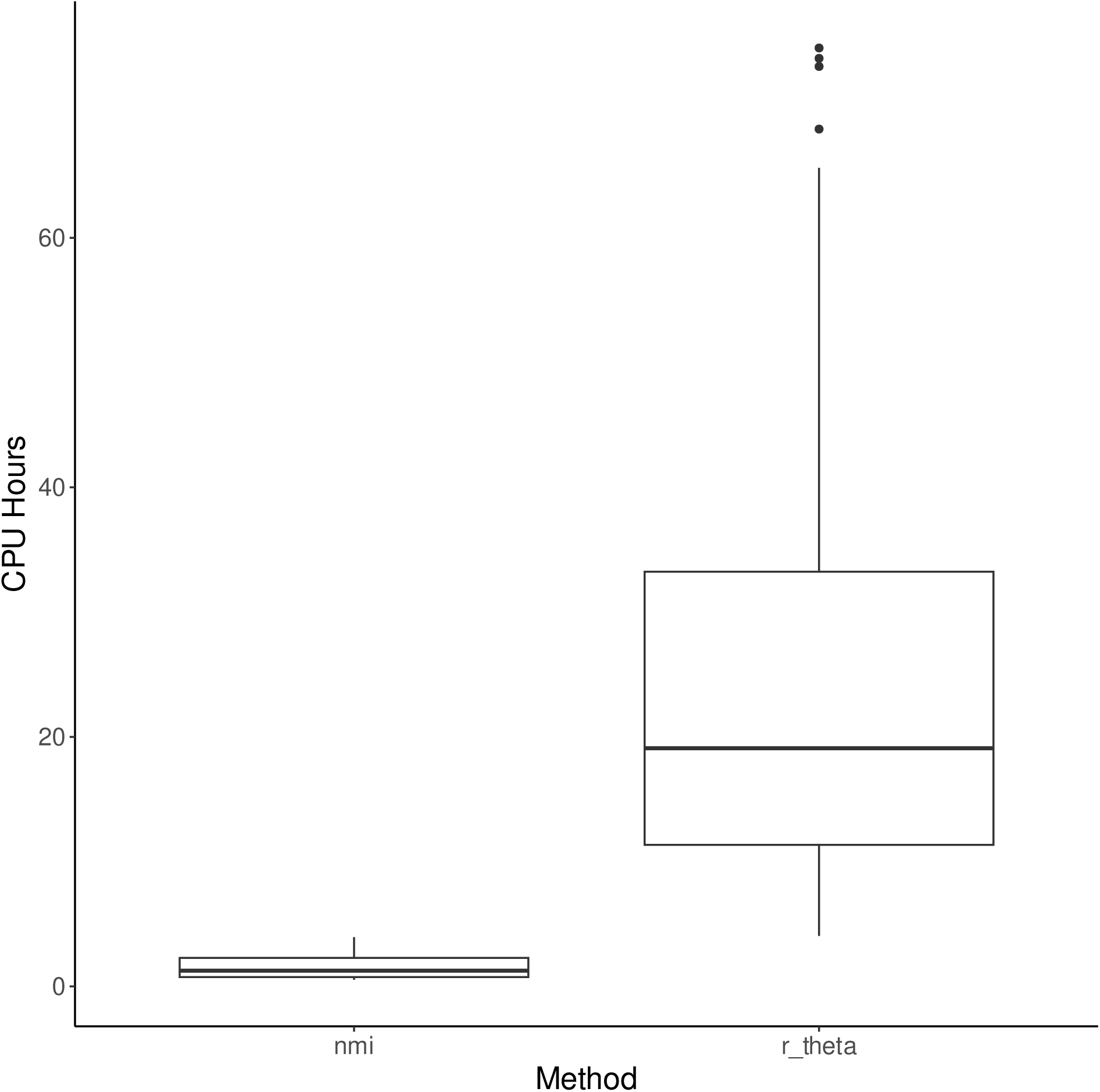
Computational savings. We calculate NMI, our proxy for genomic lability, with far less computational resource than required by traditional methods.

While time savings is an important feature of the IB, the wide range in R/theta values for *H. pylori* is emblematic of the primary weakness of most dominant recombination methods: reference bias. In Supplementary Figure 6, we show the range of R/theta values calculated for 10 randomly chosen genomes of *S aureus*, *M tuberculosis,* and *H pylori* against all possible contiguous, complete references known for these genomes. While *S aureus* and *M tuberculosis* show relatively tight ranges, *H. pylori* varies over one full unit depending on the reference chosen. Even vetted resources like RefSeq can be misleading. While compiling the data shown in Figure 6, we observed that long-branch effects from a single misclassified sequence can ruin an entire replicate. Omitting these misclassified sequences with a MASH distance-based screen clarified the relationship between NMI and R/theta (Supplemental Figure 7). We removed replicates that contained outliers in a set of all pairwise MASH distances and settled on 4.5 standard deviations from the mean to remove the heterogenous set of points near the origin while retaining the bulk of the replicates for each species. This uncertainty in RefSeq coupled with the instability inherent to reference-based analyses argues for reference free approaches like the IB. The information bottleneck is not saddled with the references required of many bioinformatic techniques including most of the dominant ones employed to detect and measure recombination.

The capacity of the bottleneck – in the Shannon sense – can be used to modulate detection. In the case of ST239 *S. aureus*, asking for just two clusters – a very narrow channel – captures more than 40% of the relevant information (Figure 5). Remarkably, these two clusters separate the core from the recombined region. Even the simplest model learns the most prominent evolutionary process. Further along the curve, fifteen clusters capture 80% of the information. To put this in perspective, the IB condenses the outsized, uninterpretable dimensions of our original data – 40 million k-mers – into just fifteen buckets while sacrificing only 20% of the information originally present in the relevance variable. Because the technique is inherently lossy, the SIB will never recover all the information originally encoded but aims to extract the most salient bits. Beyond fifteen clusters, the ST239 curve elbows, and modeling gains are slight. In this way, the relevance-compression curve defines the optimal number of clusters,^29,32^ and it is the NMI at this asymptote that best reflects the traditional R/theta.

In the light of evolution, the bend in our compression curves may have a deeper meaning. For *S. aureus*, fifteen clusters are enough to adequately capture the complete set of k-mer aggregation patterns across our chosen genomes. This point of diminishing returns may signify an opportunity for interpretive balance: not so many clusters that we drown dominant evolutionary events in noise, and not so few that we neglect to model subtle k-mer co-occurrence patterns. This particular use of the elbow method in our information theoretic context puts a crude limit on the dominant evolutionary paths taken by the elements that comprise our pangenome. And the NMI emitted at this asymptote offers a new way to describe the lability of the species.

## Software

NECK (https://github.com/narechan/neck)

## Supplemental Methods

### Information Bottleneck with neck

Neck is the program we use to execute the information bottleneck in our evolutionary context. Neck accepts two key parameters from the user: k-mer size and the number of clusters to model. The program performs a k-mer sketch of its query genomes by sliding a length *k* window across the sequence. No reference is required. We use a 64-bit hash function on canonical k-mers to generate a bottom-k sketch at the desired depth. The learning process inherent to the bottleneck procedure iterates over these k-mers, minimizing cluster distortion on each successive loop through the sketched strings. This learning process is called the sequential information bottleneck. Neck will support both assembled and unassembled sequence, and in both cases, outputs clusters of unlocalized strings. To localize strings on assembled sequence, we provide an auxiliary program called neck_paint (see below) A typical run of neck looks like this:

~~~
*neck -i genomes/ -k 19 -m 1 -c 50 -n 4 -s 10 -o out_neck_c50/*
~~~

The query genomes are found in the genomes directory *(-i genomes/*), one genome per file. In this run 19-mers are hashed at a rate of 10% *(-s 10*). In a bottom-k context this means that the lowest tenth of the hash values and their corresponding k-mers are sketched into the set of strings used in the bottleneck procedure. For this analysis, we model 50 clusters *(-c 50*) and compress the clusters over 4 iterations of the optimization (*-n 4*). Output files, including the clusters of k-mers found and the NMI value are redirected to out_neck_c50 (*-o out_neck_c50*/).

### Visualization with neck_paint

Though not strictly required, assembled genomes give a user the opportunity to contextualize clusters of k-mers generated in neck. The script neck_paint aligns k-mers from the clusters to query genomes using bowtie2 and calculates the depth of the resulting alignments with samtools. Using these depth statistics, neck_paint will calculate changepoints between regions of high and low k-mer frequency. On the core cluster, these regions typically correspond to the core genome and the recombination events. Recombination sites almost always exhibit depleted alignment in the core. De novo recombination intervals independent of any other annotations are inferred from these changepoints. If recombination regions are known, neck_paint can calculate Wilcoxon statistics on the alignment frequency differences between the known regions and the rest of the genome. We use Circos to visualize every cluster/genome pair, their associated changepoints, and any known recombination events. A typical run of neck_paint looks like this:

~~~
*neck_paint -i genomes/ -c clusters/ -l 200 -r changepoint.R -p 128 -m 3 -v 1000 -x
log.external -o out_neck_c50_paint/*
~~~

In this case, neck_paint will perform alignments, perform changepoint analysis, and generate a circos visualization for every genome *(-i genomes/*) cluster (*-c clusters/)* pair. The script changepoint.R (*-r changepoint.R*) delimits the changepoints using the PELT algorithm given the penalty parameter (*-v 1000*). The higher the penalty, the stronger the signal needed to mark a changepoint. A low penalty parameter therefore often suffers from false positives while a high penalty parameter might lack sensitivity. For the circos visualization, alignment frequency is binned into 200 basepair intervals (*-l 200*) and shown as a plot with a maximum of 3 units (*-m 3*). If available, recombination events can also be shown as a track (*-x log.external*) and the user can generate Wilcoxon statistics for non-parametric difference between frequencies within these intervals and frequencies outside of them. Because there can be many cluster/genome pairs, the process can be parallelized (*-p 128*) and the output collected into a single directory (*-o out_neck_c50_paint*).

## Supplemental Figures

**Supplemental Figure 1.**
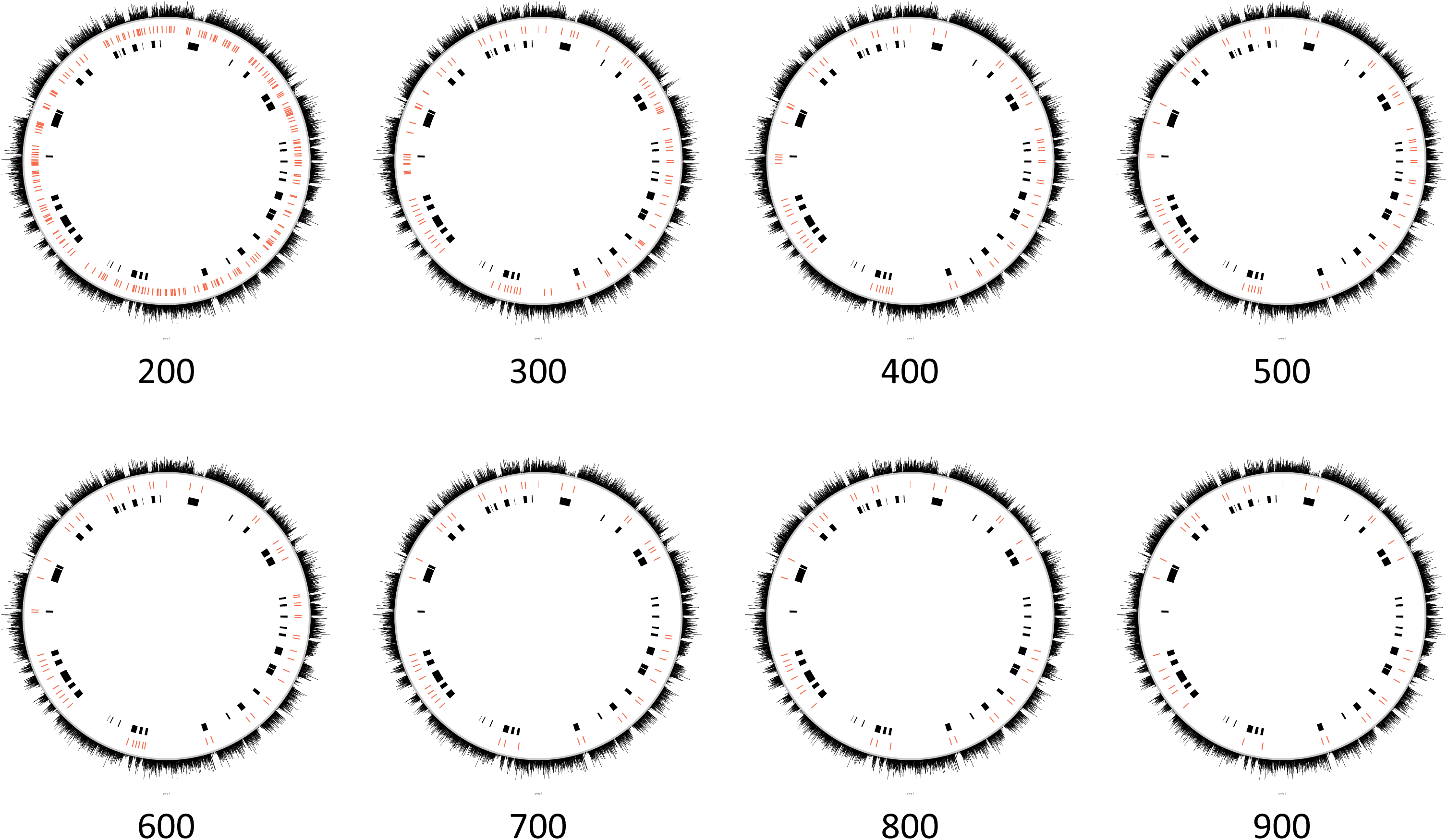
Parameter sweep in changepoint detection. Calling changepoints in the mapped k-mer frequency data is sensitive to PELT’s penalty parameter. If set too low, the analysis is mired in false postives. If set too high, we fail to call true recombination events.

**Supplemental Figure 2.**
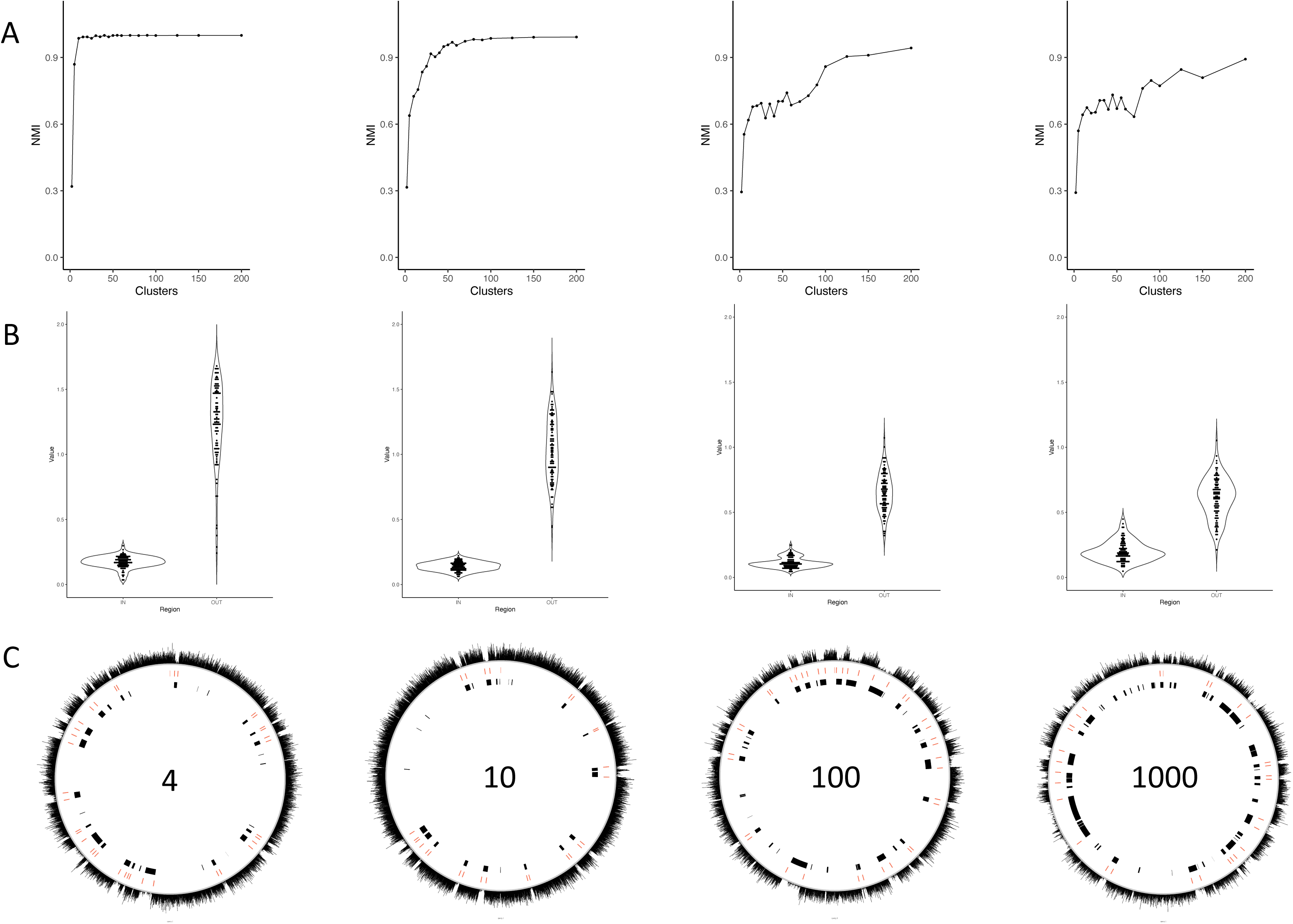
Compression efficiency with large datasets. We simulated recombination in 100 sets of 4, 10, 100, and 1000 one megabase genomes. In A we show the rate of compression efficiency with increased modeling in the example relevance-compression curves. More complex datasets require more clusters to capture comparable amounts of information. In B we show that compression noise in more complex datasets is due to the narrowing gap between core cluster k-mers captured outside versus inside simulated recombination events. In C, we show examples of k-mers from the core IB cluster mapped to one of the genomes. Simulated events are displayed in black, and changepoints are shown as red tick marks delimiting significant changes in the frequency of the k-mer distribution.

Supplemental Figure 2. Simulation across various rates of recombination and levels of modeling. NMI decreases with increasing recombination at approximately the same rate across different levels of modeling. Modeling more clusters, captures more information. For lower levels of recombination, over-modeling may cause a decrease in useful information captured.

**Supplemental Figure 3.**
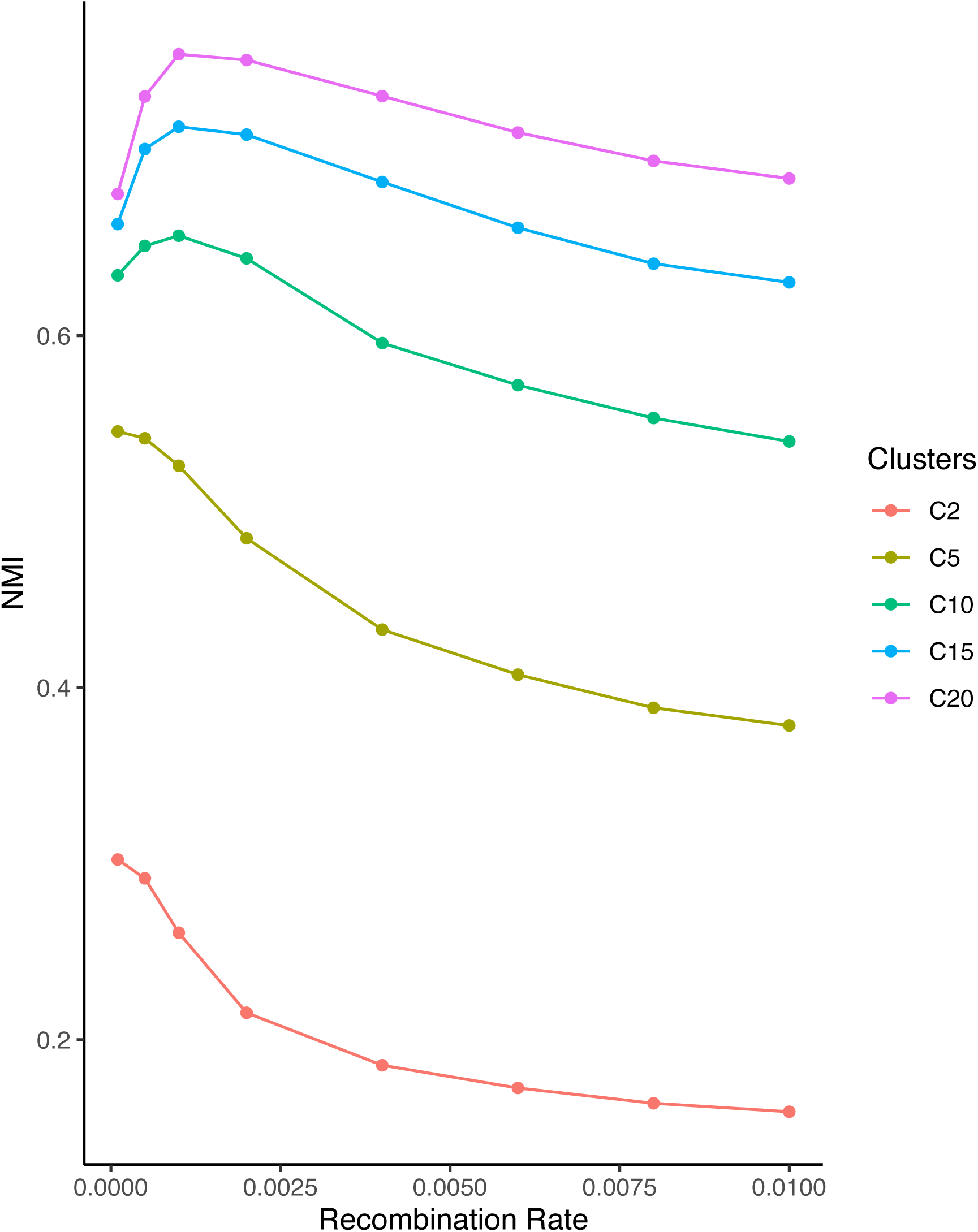
Relevance compression curves across 28 species. We show how increasing the number of clusters modeled, increases the amount of information captured across the 28 most sequenced microbial species in GenBank.

**Supplemental Figure 4.**
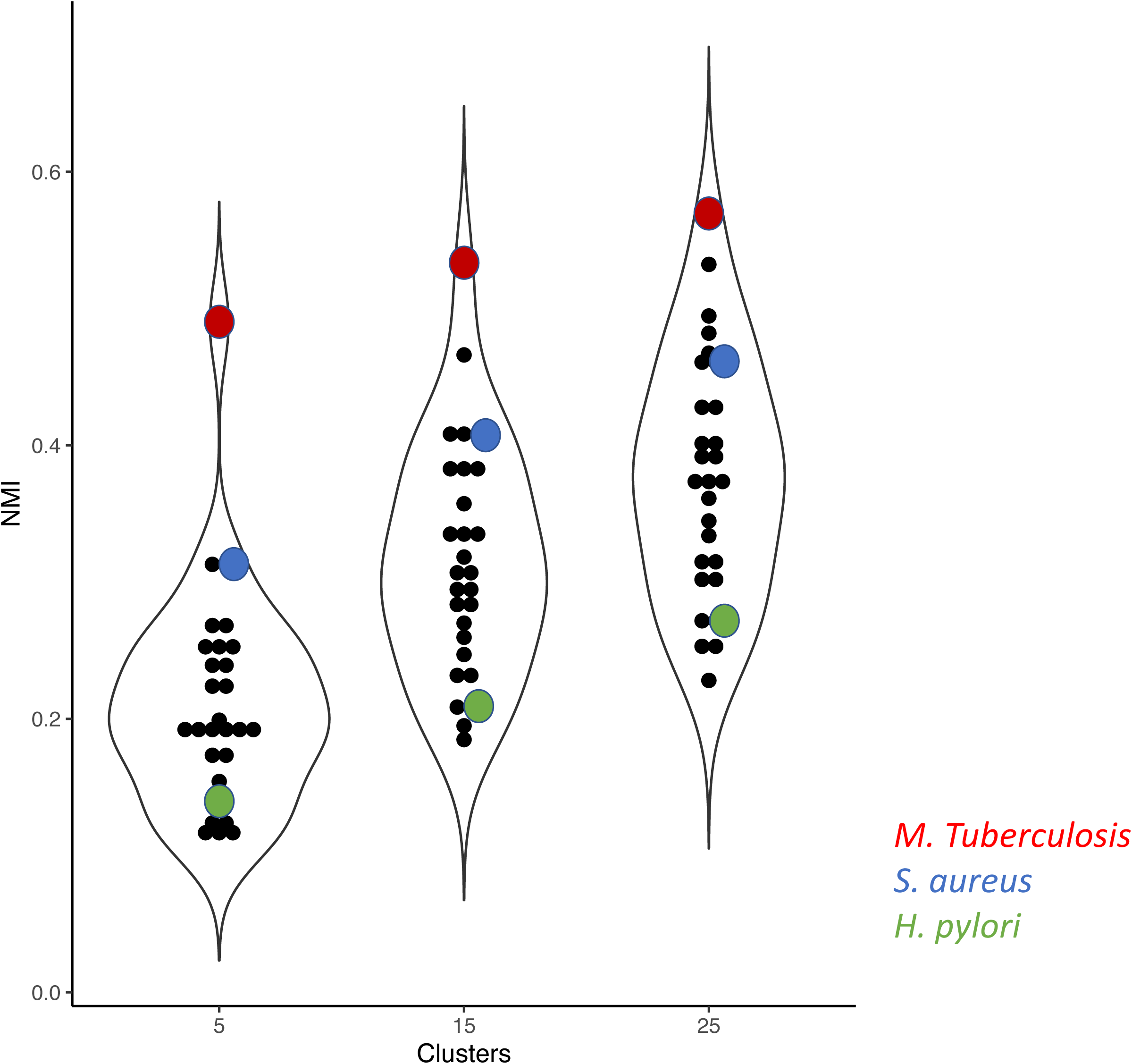
Species-level optimal compression. Regardless of species, the NMI stabilizes at about 50 randomly selected genomes. Increased genomic information does not seem to add greater complexity beyond that point.

**Supplemental Figure 5.**
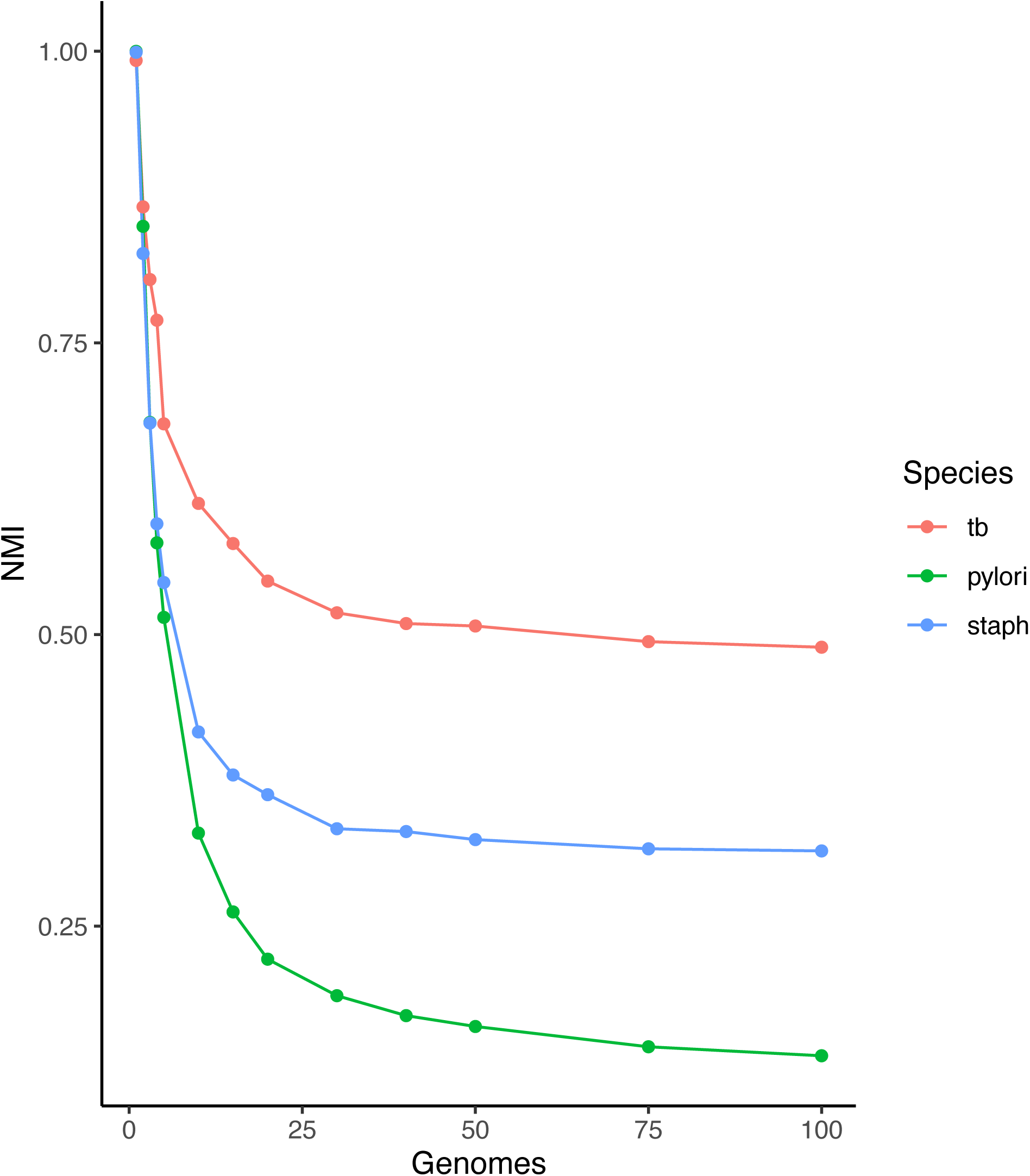
Reference bias in traditional methods. We show that for *H. pylori*, R/theta varies over a large range depending on the reference chosen.

**Supplemental Figure 6.**
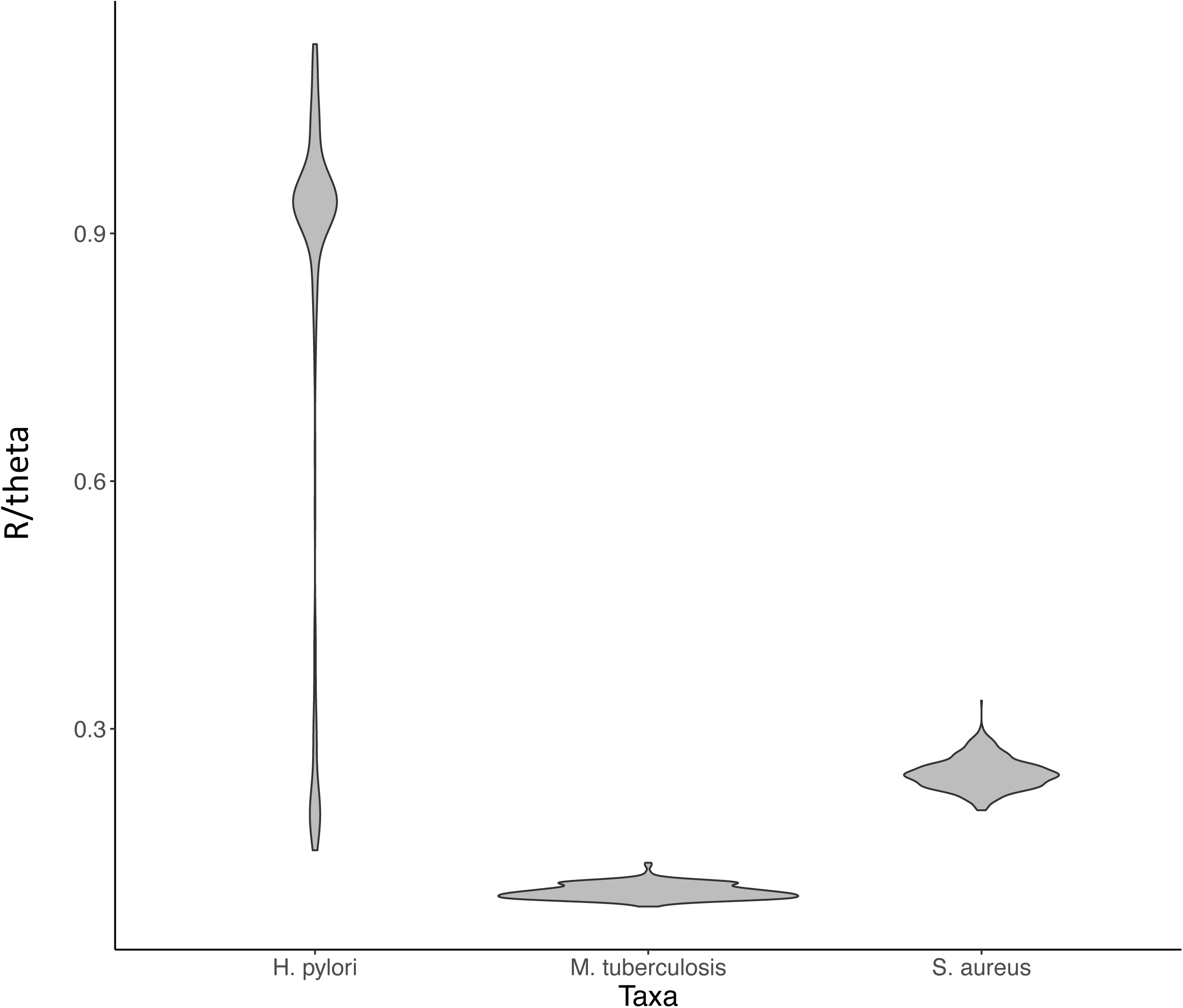
The effect of increased stringency in outlier removal. RefSeq contains misannotated sequences that cause severe long branch effects. We screened any replicates with these sequences using an all against all MASH distance analysis. We show the change in the replicates kept as the stringency for removing experiments is increased from three standard deviations of the mean to 4.5 standard deviations. In the panel with all replicates, we show that the points removed cluster close to the origin.

**Supplemental Figure 7.**
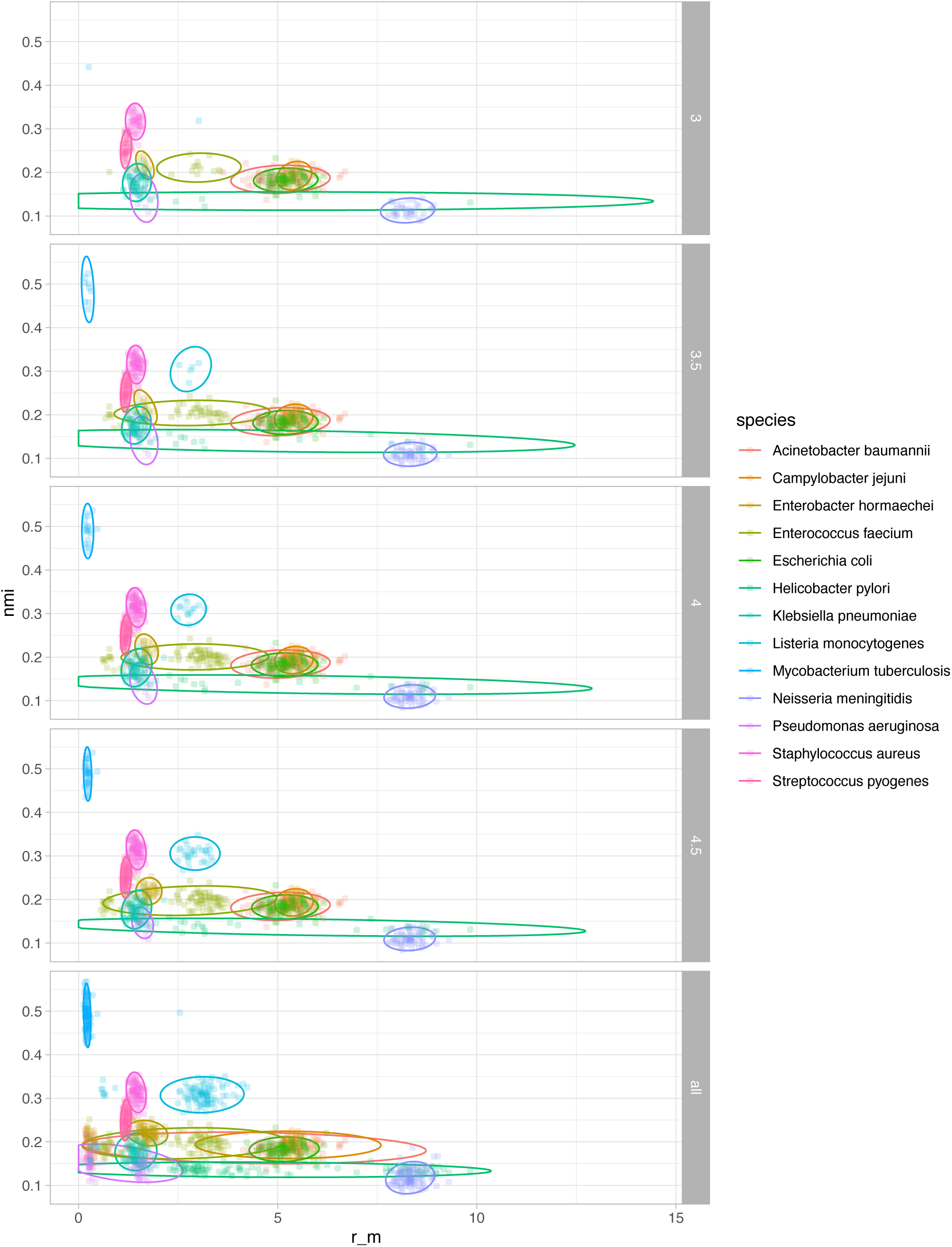
MASH screen of long branches. Misclassified sequences in RefSeq can cause estimates for r/m to vary widely. We screened replicates that contained outliers in the set of all pairwise MASH distances. Shown are variants of Figure 6B with successively more replicates removed as the threshold for removal is increased. At 4.5 standard deviations from the mean, the cluster of heterogenous points near the origin disappears while retaining most of the replicates for each species.

## Notes

### Competing Interest Statement

The authors have declared no competing interest.

### Summary of Updates

We added significant emphasis on Normalized Mutual Information as a proxy for a species tendency to recombine.

